# Female factors are important for the seminal Sex Peptide’s association with sperm in mated D. melanogaster

**DOI:** 10.1101/2022.05.13.491855

**Authors:** Snigdha Misra, Akanksha Singh, Mariana F. Wolfner

## Abstract

**Background:** Male-derived seminal fluid proteins (SFPs) enter and induce a myriad of physiological and behavioral changes in mated female flies optimizing fertility. Many post-mating changes in female Drosophila melanogaster persist for ∼10-14 days, because the seminal protein that induces them, Sex Peptide (SP), is retained long-term in females by binding to sperm, with gradual release of its active domain from sperm. Several other “long-term response SFPs” (LTR-SFPs) “prime” sperm to bind SP. Whether female factors play a role in this process is unknown, though it is important to study both sexes for a comprehensive physiological understanding of this reproductive process and for consideration in models of sexual conflict.

**Results:** We report here that sperm in male ejaculates bind SP more weakly than sperm that have entered females, and that the amount of SP and other SFPs bound to sperm increases with time and transit of individual seminal proteins within the female reproductive tract. Thus, female contributions are needed for maximal and appropriate binding of SP, and other SFPs, to sperm. Towards understanding the molecular roles of the female, we found no dramatic change in pH of the female reproductive tract after mating and that initial binding of SP to sperm is normal in females with ablated or defective spermathecal secretory cells and/or parovaria, but that higher levels of SP (and sperm) are retained in the latter females.

**Conclusion:** This study reveals that the SP pathway is not entirely male-biased and that females also contribute to regulating it.

## Background

Molecular interactions between the male’s seminal proteins (SFPs), sperm, and the female’s reproductive tract (FRT) are fundamental to successful reproduction [1–4]. For example, in Drosophila melanogaster, SFPs derived from glandular tissues of the male’s reproductive tract induce ovulation (ovulin [5]) or participate in the formation of the mating plug (Acp36DE, pEBme, pEBII; [6–8]) within the reproductive tract, as well having a variety of other systemic effects [9–13]. In addition, several Drosophila SFPs associate with sperm [4,14,15]. One sperm-associated SFP, Sex Peptide (SP), is retained in the female long-term due to its association with sperm [16]. SP’s active C-terminal region is gradually released from sperm by trypsin cleavage, and induces long-term post-mating responses such as increased egg production and decreased receptivity to remating [17–20].

Previous studies have shown that SP’s binding to sperm requires several other SFPs, acting in a network (the “long-term response (LTR) network” [3,4]). Two SFPs in this network (Seminase, CG17575; [21,22]) facilitate the binding of other SFPs to sperm, but do not themselves bind sperm. Other SFPs in the network (CG1656 [lectin-46Ca], CG1652 [lectin-46Cb], CG9997, Antares; [14,15,23]) bind to sperm transiently; their action is thought to “prime” sperm to retain SP. However, thus far all molecules known to promote SP binding to sperm have been male-derived SFPs. Although some female proteins, such as Fra mauro, Hadley, Esp, and Sex Peptide Receptor (SPR) are known to be necessary for SP-induced post-mating responses, their action is downstream of the binding of SP to sperm [24–28].

It was unknown whether the female also contributes to the binding of SP to sperm, but several recent findings suggest the importance of testing this possibility. First, the molecular composition of sperm changes within the mated female, by the association of multiple female-derived proteins with sperm ([29–32]); such female proteins would be in a position to affect SP’s binding to sperm. Second, female molecules play roles in modification (cleavage) of some SFPs in D. melanogaster [33], and in the proteolytic dissolution of the mating plug in cabbage-white butterflies [34], again indicating that FRT proteins can have direct effects on SFPs and their molecular milieu. Third, active involvement by females in relative paternity proportions following mating with two males suggests that female molecules or cells can interact with ejaculate components (at least, sperm) [35–37].

Therefore, here we tested whether female contributions affect the binding of SP, and other SFPs, to sperm. We found that levels of sperm-bound SFPs are weak or undetectable in the male ejaculate, but sperm-binding by SFPs, including SP, becomes detectable (or increases) after ejaculate enters the female. The pattern and the signal intensity of binding of individual SFPs to sperm differ temporally and spatially within the FRT. This increase in their signal intensity level indicates that female components must play a role in priming sperm to bind SP. Our investigations took two different approaches to identify the nature or source of female molecules that facilitate SFP-sperm binding. First, we found no dramatic change in the pH of the female reproductive tract after mating, in contrast to the situation in humans, but similar to what has been reported for other non-human mammalian models such as mice [38]. Second, upon disabling two secretory tissues in the FRT (spermathecal secretory cells (SSCs) and parovaria; [39–41]) we observed no dramatic effects on initial binding of SP to sperm. However, loss of secretions from SSCs resulted in the retention of sperm-associated SP, long term (4d after the start of mating (ASM)).

Our finding that females, as well as males, contribute molecules needed to bind SFPs to sperm and to cleave SP’s active region from sperm has implications for understanding the molecular cooperation between the sexes that leads to optimal fertility, as well as for models of sexual conflict, and motivates future studies to identify these female players.

## Results

### 1. Sex Peptide binds sperm weakly in the male ejaculate but its binding increases within the female bursa and seminal receptacle

To test whether female factor(s) (molecule(s) or environment) affect the binding of SP to sperm, we compared the signal intensity of anti-SP staining on sperm before (in male ejaculate) and after mating (in female bursa [uterus] and seminal receptacle). We reasoned that if the signal intensity in the male ejaculate did not change after mating, this would mean that components of the male ejaculate are sufficient to fully facilitate SP-sperm binding without requiring female factor(s).

We isolated sperm from ejaculates exuded by males (Eja; 0 min), sperm in the mated female’s bursa (35 min after the start of mating; ASM) or stored in her seminal receptacle (SR; 2 hrs ASM). The amount of SP bound to sperm was determined by quantifying the signal intensity of the immunofluorescence of anti-SP along the sperm tail in all three samples. The signal intensity for SP detected on sperm was weakest in male ejaculates (Fig. 1A-A’ and I; Mean±SE=2.1±0.71 AU (AU=arbitrary units)). It was higher in sperm isolated from mated female bursae. Sperm isolated from mated female bursae (35 min ASM) had a “spotty” pattern of anti-SP staining, with anti-SP immunofluorescence appearing in bright and dim specks all along sperm (Fig. 1C-C’ and I, Mean±SE=4.67±0.71 AU; compare to Fig. 1A-A’). This suggested that although the quantity of SP bound to sperm increased in the female’s bursa, sperm were not uniformly saturated with SP. Sperm isolated from the seminal receptacles (2 hrs ASM) had the highest signal intensity of SP, suggesting that the amount of SP detected on sperm was greatest in the sperm storage organ. Staining for SP on sperm isolated from seminal receptacles was consistent and uniform along the sperm (Fig. 1E-E’ and I; Mean±SE=13.15±0.71 AU), similar to what has been reported previously [14–16]. Since the amount of SP detected on sperm gradually increases after they enter the FRT, our results suggest a possible role of female factor(s) in assisting SP binding to sperm.

**Figure 1.**
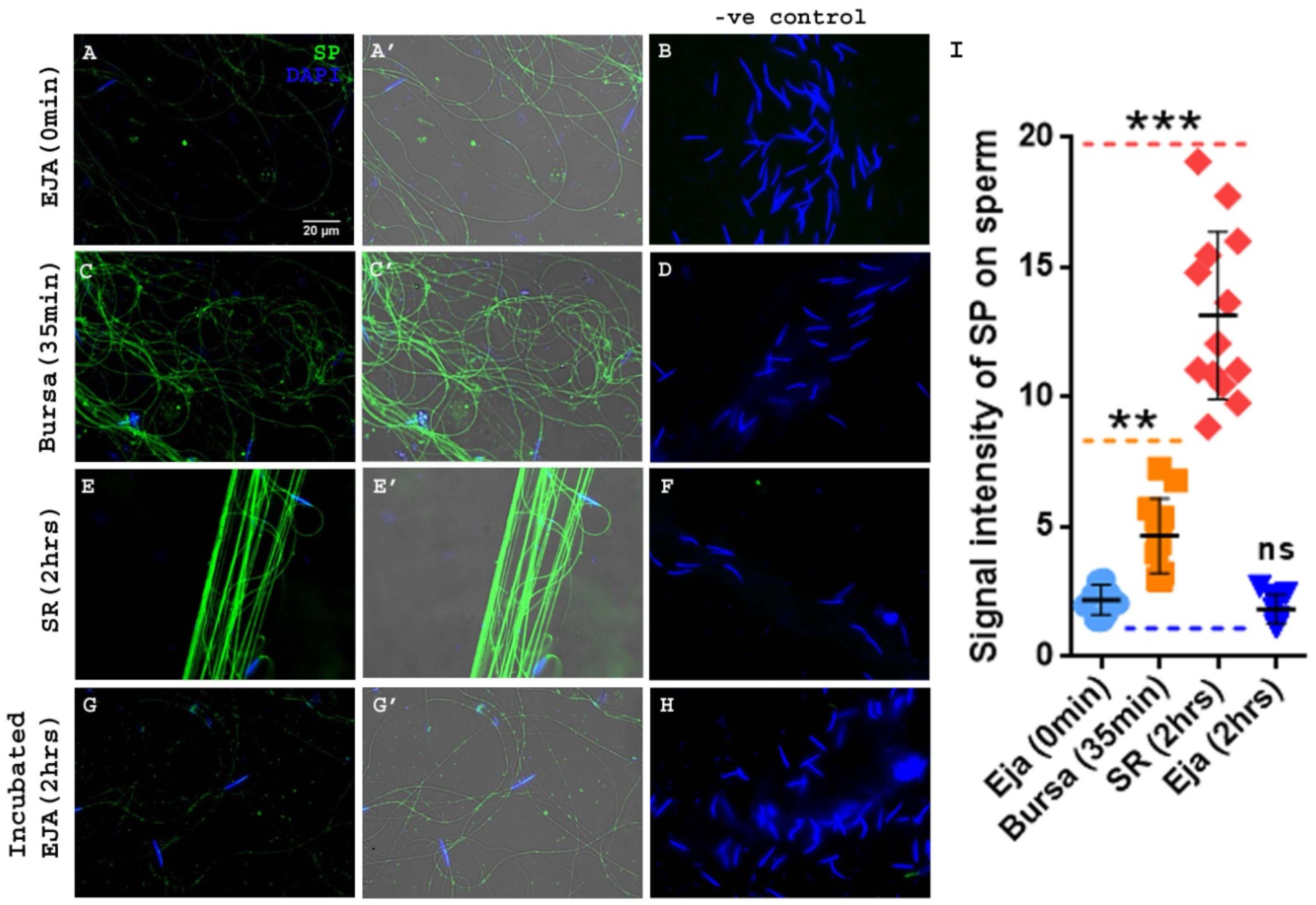
Amount of SP detected on sperm gradually increases from male ejaculate to female bursa, and is highest in seminal receptacles. Pre-mating ejaculate samples were collected from Fru>dTRPA1 males exposed to high temperature [33]. Post mating sperm samples were isolated from wild type (CS) females that had mated to wildtype (CS) males and were frozen at 35 min and 2 hrs ASM. Sperm heads were stained with DAPI (blue) and anti-SP staining was visualized with Alexa fluor 488, staining the sperm tail (green) and sperm head (cyan; overlapping blue/green). **(A-A’)** Sperm isolated from ejaculate exuded by Fru>dTRPA1 males. **(C-C’)** Sperm isolated from bursa of the wildtype mated female, frozen at 35 min ASM. **(E-E’)** Sperm isolated from the seminal receptacle of the wildtype mated female, frozen at 2hrs ASM. **(G-G’)** Sperm isolated from Fru>dTRPA1 male’s ejaculate that was kept incubated for 2 hrs in 1X PBS, after exudation. **B, D, F and H** are negative controls for A, C, E, and G panels, with only secondary antibody (anti-rabbit, Alexa fluor 488) and no primary antibody (anti-SP) incubation. **A’, C’, E’ and G’** panels have an additional transmitted light channel overlay to highlight the outline of sperm tail specifically in samples where the staining was very weak (ejaculate). **(I)** Relative signal intensity of SP on sperm at three different stages and time points, p***<0.001, p**<0.01, ns=not significant, error bars show Mean±SE AU (n=10; Bar = 20µm; AU stands for arbitrary units).

Several SFPs (proteases, prohormones, and others) either mediate or undergo post-mating modifications en route to or after transfer to the FRT [5,33,42], some of which are crucial for inducing or maintaining post-mating responses in mated females. We thus wondered whether the gradual increase in amount of SP detected on sperm within the FRT is because of a need for time for the male molecules to undergo requisite modifications.

The intensity of SP signals on sperm was observed to be highest in sperm isolated from the seminal receptacle at 2 hrs ASM, suggesting that this is the maximum time that would be required by the male molecules to act (or to undergo any necessary modifications). To test if time alone is sufficient to maximize SP’s binding to sperm, we collected ejaculates exuded from males and incubated them for 2hrs in 1X PBS before processing them for anti-SP staining. We did not observe any change in the signal intensity or distribution of anti-SP on sperm (Fig. 1G-G’ and I, Mean±SE=1.84±0.71 AU) in incubated ejaculates relative to signals on sperm isolated from un-incubated ejaculates (Fig. 1A-A’, Eja 0 min).

This suggested that time alone is not sufficient to maximize SP’s binding to sperm. Thus, female factor(s) likely contribute to, or facilitate, SP-sperm binding.

### 2. LTR-SFPs bind to sperm in the male’s ejaculate or mated females with patterns or timing different from those of SP

Given LTR-SFPs’ role in SP’s sperm-binding, we wondered whether the pattern of sperm-associated CG1656, CG1652, CG9997, and Antares (Antr) in sperm isolated from three different sites/times used above paralleled that of SP. We examined the presence of bound LTR-SFPs to sperm by experiments analogous to those shown in Fig. 1 for SP, using sperm isolated from the male’s ejaculate (0 min after exudation), mated female’s bursa (35 min ASM), and seminal receptacle (2 hrs ASM).

We observed a lower signal intensity for CG1656 (Fig. 2A and Fig. 2C’; Mean±SE=6.87±0.79 AU) and Antr (Fig. 2D and Fig. 2F’; Mean±SE=6.97±0.1.22 AU) on sperm in ejaculate compared to that on sperm inside the female (Figs. 2B (11.74±0.79 AU), C (11.83±0.77 AU) and C’ for CG1656 and Figs. E (14.67±1.26 AU), F (15.22±1.34 AU) and F’ for Antr). However, the signal intensity of sperm-bound CG1656 and Antr did not differ between sperm isolated from the bursa (Fig. 2B, E) vs. those from the seminal receptacles (Fig. 2C, F). This suggests that although the amount of these LTR-SFPs bound to sperm increases post-mating, their maximal binding had already occurred in the bursa of the mated female, in contrast to SP whose sperm-binding reached its highest levels in the female’s seminal receptacle.

**Figure 2.**
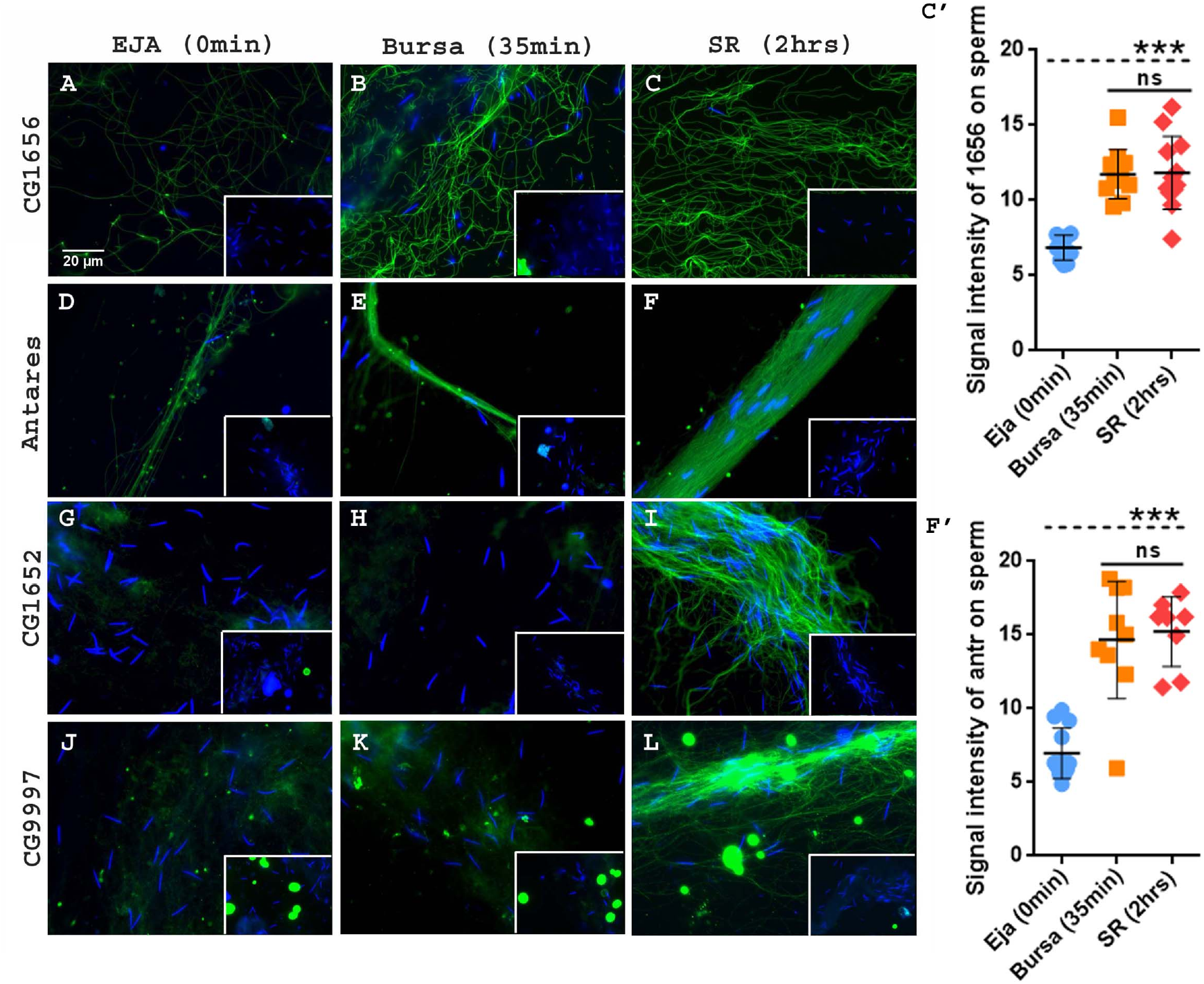
The levels of LTR-SFPs bound to sperm increase from male ejaculate to sperm stored in female seminal receptacle, but the pattern differs from SP’s. Pre-mating ejaculate samples were collected from Fru>dTRPA1 males exposed to high temperature. Post mating sperm samples were isolated from wild type (CS) females that had mated to wildtype (CS) males and frozen at 35 min and 2 hrs ASM. Sperm heads were stained with DAPI (blue) and LTR-SFPs were visualized with Alexa fluor 488, staining the sperm (green). Sperm isolated from male ejaculate immediately after exudation **(A, D, G and J)** were probed for CG1656, Antares, CG1652 and CG9997, respectively. Sperm isolated from mated female bursa, frozen at 35 min ASM **(B, E, H and K)** were probed for CG1656, Antares, CG1652 and CG9997, respectively. Sperm isolated from mated female’s seminal receptacle, frozen at 2hrs ASM **(C, F, I and L)** were probed for CG1656, Antares, CG1652 and CG9997, respectively. The insets show the negative controls for their respective panels. Sperm samples in negative controls were incubated with only secondary antibody (anti-rabbit, Alexa fluor 488) but no primary antibody (anti-LTR-SFP) incubation, as mentioned previously. **(C’ and F’)** show relative signal intensity of CG1656 and Antares on sperm at three different stages and time points, p***<0.001, ns=not significant, error bars show Mean±SE AU (n=10; Bar = 20µm; AU stands for arbitrary units).

CG1652 and CG9997 differed in their sperm-binding pattern from CG1656 and Antr. We could not detect binding by either CG1652 or CG9997 to sperm in the ejaculate (Fig. 2G, 2J respectively) or in the bursa of the mated female (Fig. 2H, 2K respectively). However, we saw a strong signal for both proteins in sperm isolated from seminal receptacles of mated females (Fig. 2I, 2L respectively), consistent with our previous report that these proteins are bound to sperm in seminal receptacles [4,14,21]. The regions of association and distribution that we observed for these SFPs (SP, CG9997, and Antr on the head and tail of stored sperm; CG1652 and CG1656 detectable only on the tail of stored sperm) were also consistent with previous reports [14,21].

We also assessed the two LTR-SFPs, Seminase and CG17575, that had previously been reported to not bind to stored sperm [21,22]; in addition to confirming that finding, our experiments showed that these two SFPs exhibit no sperm-binding in the ejaculate either (ejaculate: Fig. 2S A and D; mated female’s bursa: Fig. 2S B and E; seminal receptacle: Fig. 2S C and F).

Thus, the binding patterns/timing of LTR-SFPs differed from those of SP, and fall into three groups: (1) CG1656 and Antr, which bind to sperm in the ejaculate, increase their binding once inside the female, but do not show the additional increase in binding in the seminal receptacle, as was seen for SP; (2) CG1652 and CG9997 show no detectable binding to sperm until they are inside the female’s seminal receptacle; (3) Seminase and CG17575 show no detectable binding to sperm.

### 3. Membrane-associated pH sensors indicate that the pH of the virgin or mated female reproductive tract falls in the range of 6-7.4

The extracellular environment in the FRT can have important effects on sperm (reviewed in [31]). For example, factors such as the pH and viscosity of FRT fluids regulate the storage, viability, and/or motility of mammalian sperm [38,43]. The small size and convoluted shape of the D. melanogaster FRT made it impossible to measure its pH by conventional means, so we used genetically encoded pH sensors to estimate whether there are any major changes in FRT pH following mating, with the caveat that our results are limited to the sensitivity of these sensors. To monitor potential pH changes within the FRT before and after mating, we expressed membrane-associated pH sensors (MApHS; [44,45]) in particular regions of the FRT. We first expressed an ecliptic pHluorin sensor (pHluorinE) fused to an extracellular domain. The fusion protein also contains tdTomato (tdTom) in its intracellular domain. pHluorinE is brightest at pH 7.4 under 475nm excitation but gets dimmer as the pH drops, and loses fluorescence at pHs below 6. Thus, for example, if tissues of the virgin FRT had a pH in the acidic range (e.g. pH<6; as observed in humans) but spiked up in pH to pH≥7.4 following mating, we would expect that pHluorinE should exhibit no green fluorescence in FRTs of unmated females (pH<6) while still retaining red fluorescence of tdTom, whereas in newly mated FRTs (pH≥7.4) that same sensor would fluoresce both green (pHluorinE) and red (tdTom).

Because ubiquitous expression of pHluorinE was lethal, we drove its expression in tissues of the FRT using tissue-specific Gal4 drivers. We used Oamb-Gal4 [46] and Send1-Gal4 [39] to drive the expression of pHluorinE in the oviduct and spermathecal secretory cells (SSCs) of the FRT, respectively. The Oamb>pHluorinE and Send1>pHluorinE females were dissected and their reproductive tracts were imaged for pHluorin signals, before and after mating (35min ASM).

We observed overlapping pHluorinE (green) and tdTom (red) signals in the oviduct (yellow) of Oamb>pHluorinE virgin females (Fig. 3A). The signals were also observed in the parovaria (aka female accessory glands) of these females, though the fluorescence of pHluorinE was lower than observed in the oviduct. Mated Oamb>pHluorinE females dissected at 35min ASM showed a similar pattern of pHluorinE and tdTom signals in their oviducts and parovaria (Fig. 3B). Control females (Oamb>CyO) did not show red or green fluorescence in their reproductive tracts, as expected (Fig. 3C and D).

**Figure 3.**
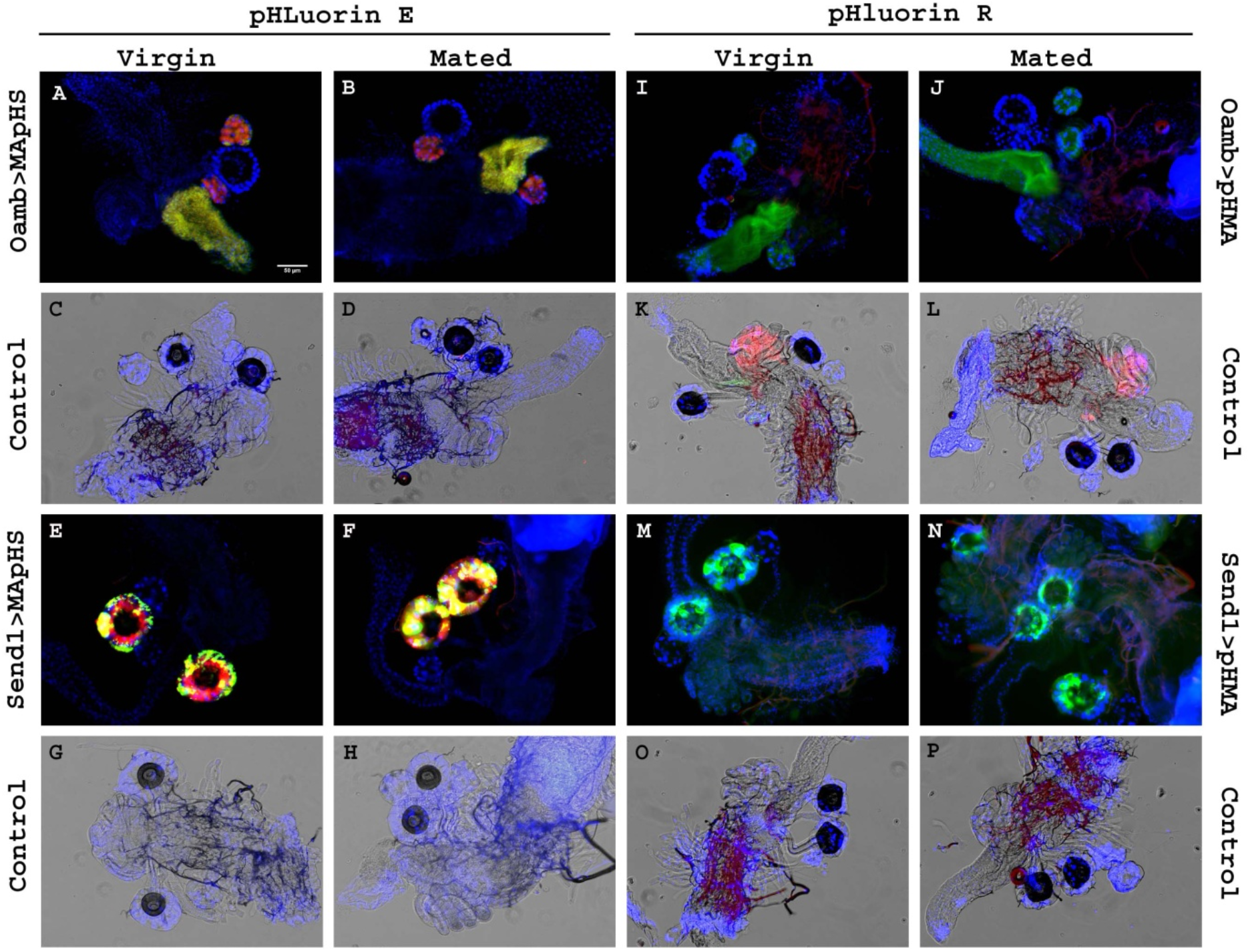
Membrane-associated ecliptic (MApHS) and ratiometric pH sensors (pHMA) indicated that the pH of the female reproductive tract is in the range of 6.0-7.4, with no drastic shift as a result of mating. Expression of membrane associated pHluorin E (MApHS) by Oamb Gal4 in the oviduct of virgin **(A)** and mated **(B)** females. Oamb>CyO Controls for Oamb>MApHS, virgin **(C)** and mated **(D)**. Expression of pHluorin E (MApHS) by Send1-Gal4 [39] in the SSCs of virgin **(E)** and mated **(F)** females. Send1>CyO Controls for Send1>MApHS, virgin **(G)** and mated **(H)**. pHluorin E (green) and tdTom (red) signals evident as overlapping yellow signals in oviduct and SSCs. Expression of pHluorin R (pHMA) by Oamb-Gal4 [46] in the oviduct of virgin **(I)** and mated **(J)** females. Controls for Oamb>pHMA, virgin **(K)** and mated **(L)**. Expression of pHluorin R (pHMA) by Send1 Gal4 in the SSCs of virgin **(M)** and mated **(N)** females. Send1>CyO Controls for Send1>pHMA, virgin **(O)** and mated **(P)**. pHluorin R (green) in oviduct and SSCs. Nuclei of the female RT stained with DAPI (blue). (n=10; Bar = 50µm).

We also observed overlapping pHluorinE (green) and tdTom (red) signals in the spermathecal secretory cells (SSCs; yellow) surrounding the spermathecal cap in Send1> pHluorinE virgin females (Fig 3E), and no difference in the pattern of expression or strength of the signal in mated Send1>pHluorinE females (Fig 3F). Control females (Send1>CyO) did not give any fluorescence signals in SSCs, as expected (Fig. 3G and H). Our data indicate that different sites or tissues within the female reproductive tract have pHs between 6.0 to 7.4 and that there is no change in the pH of the female reproductive tract outside of this range post-mating.

We also used a pH-sensitive ratiometric derivative of GFP, pHluorinR, to further examine the pH in virgin and mated FRTs. This indicator is a fusion of pHluorin and the myosin actin-binding domain (pHMA; [44,47]). It fluoresces at 470 nm (green) at neutral pH (7.4), but upon acidification (pH<6), pHluorinR exhibits a spectral shift and fluoresces at 415 nm (red). We drove the expression of pHluorinR in different tissues of the female reproductive tract. Ubiquitous expression of pHluorinR was lethal, so we again used Oamb-Gal4 and Send1-Gal4 to drive the expression of pHluorinR in the female reproductive tract.

We observed pHluorinR signals (green) in the oviduct of Oamb>pHluorinR virgin females (Fig. 3I). We did not observe any spectral shift from green to red in the oviducts of mated Oamb>pHluorinR females: at 35min ASM, the intensity of the green signal was the same as observed in virgins (Fig 3J). Control females (virgin or mated) did not give any fluorescence signals in the reproductive tract (Fig. 3K and L). Similarly, we observed a green signal in the spermathecal secretory cells (SSCs) surrounding the spermathecal cap in Send1>pHluorinR virgin females (Fig 3M) and no shift in fluorescence from green to red in mated females (Fig 3N). Control females gave no fluorescence signals in SSCs, as expected (Fig. 3O and P). These results confirmed the pH of female reproductive tissues to be in the range of 6.0 to 7.4 and also confirmed that, to the limit of sensitivity of available sensors, there is no detectable shift in the pH outside of this range, post-mating.

### 4. Ablation of spermathecal secretory cells (SSCs) in the female reproductive tract does not affect the initial binding of SP or LTR-SFPs to sperm

To test whether the increase in the signal intensity of SFPs bound to sperm once inside the FRT is the consequence of secretions from FRT tissues, we examined the effect of loss of female reproductive tract secretions on the intensity and timing of SFP binding to sperm. The secretions from SSCs are known to regulate the storage and motility of sperm in sperm storage organs [39–41]. We ablated the SSCs that line the spermathecal cap by driving the expression of misfolded protein Rh^1G69D^ [48,49] in these cells (Fig. 4A-D) or by using Hr39 mutants (Please see supplementary material: Text (1) and Fig. 5S1). Hr39 is needed for the formation of secretory units in the female reproductive tract; Hr39 mutants have been reported to exhibit defective (ablated) SSCs and parovaria [40,41]. Although we were not able to completely ablate all SSCs by either method, we examined if SP-sperm binding was abnormal in these SSC deficient mutant females.

**Figure 4.**
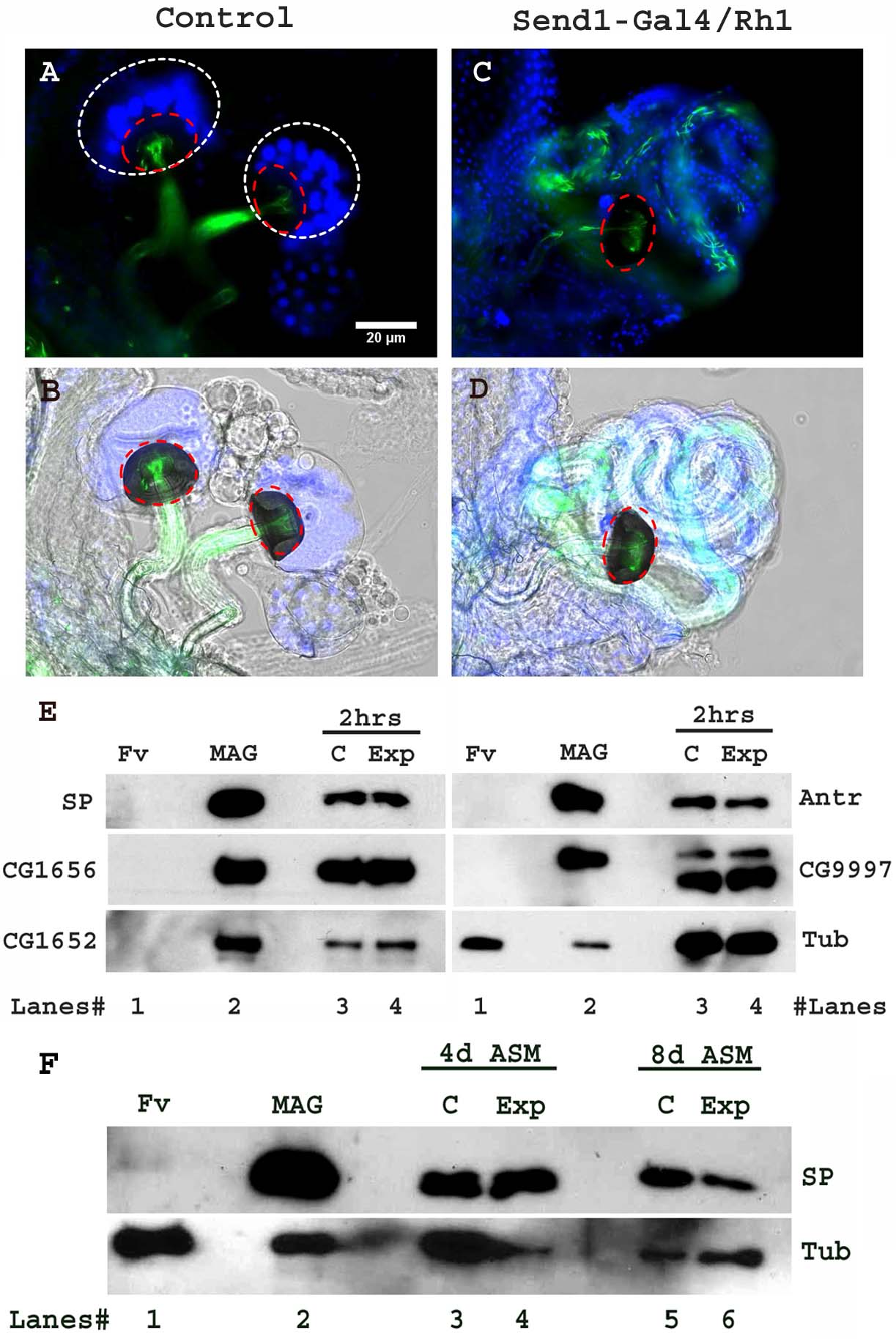
Ablated SSCs in Send1> Rh1 females affects neither the binding of SP to sperm (at 2hrs ASM) nor the long-term gradual cleavage of SP from sperm (at 4d or 8d ASM). Expression of ER stress-inducing Rh1^G69D^ [48] by spermathecae-specific Send1-Gal4 results in ablation of SSCs. **(A-B)** Control (Send1>CyO) females mated with ProtB-eGFP males (eGFP-tagged sperm; green) show normal numbers of SSCs (stained with DAPI; white dotted circle) lining the spermathecal cap (red dotted circle). **(C-D)** Experimental (Send1>Rh1^G69D^) females mated with ProtB-eGFP males show ablated SSCs (stained with DAPI) lining the spermathecal cap (red dotted circle); n=10; Bar = 20µm. **(E)** Western blot probed for SP and indicated LTR-SFPs at 2hrs ASM. **Lanes# 1: Fv**, reproductive tract (RT) of 4 virgin females (negative control), **2: MAG**, 1 pair of male accessory glands (positive control), **3: C**, sperm dissected from SR of 30 control (Send1>CyO) females mated to wild type (CS) males at 2hr ASM, **4: Exp**, sperm dissected from SR of 30 experimental (Send1>Rh1^G69D^) females mated to wild type (CS) males at 2hr ASM. Lanes were probed for SP and LTR-SFPs CG1656, CG1652, Antares and CG9997 as described in the text. **(F)** Western blot probed for SP at 4 and 8 days ASM **Lanes# 1: Fv**, reproductive tract (RT) of 4 virgin females (negative control), **2: MAG**, 1 pair of male accessory glands (positive control), **3: C**, sperm dissected from SR of 30 control (Send1>CyO) females mated to wild type (CS) males at 4 days ASM, **4: Exp**, sperm dissected from SR of 30 experimental (Send1>Rh1^G69D^) females mated to wild type (CS) males at 4 days ASM, **5: C**, sperm dissected from SR of 30 control (Send1>CyO) females mated to wild type (CS) males at 8 days ASM, **6: Exp**, sperm dissected from SR of 30 experimental (Send1>Rh1^G69D^) females mated to wild type (CS) males at 8 days ASM. Lanes were probed for SP. Tubulin served as loading control.

**Figure 5.**
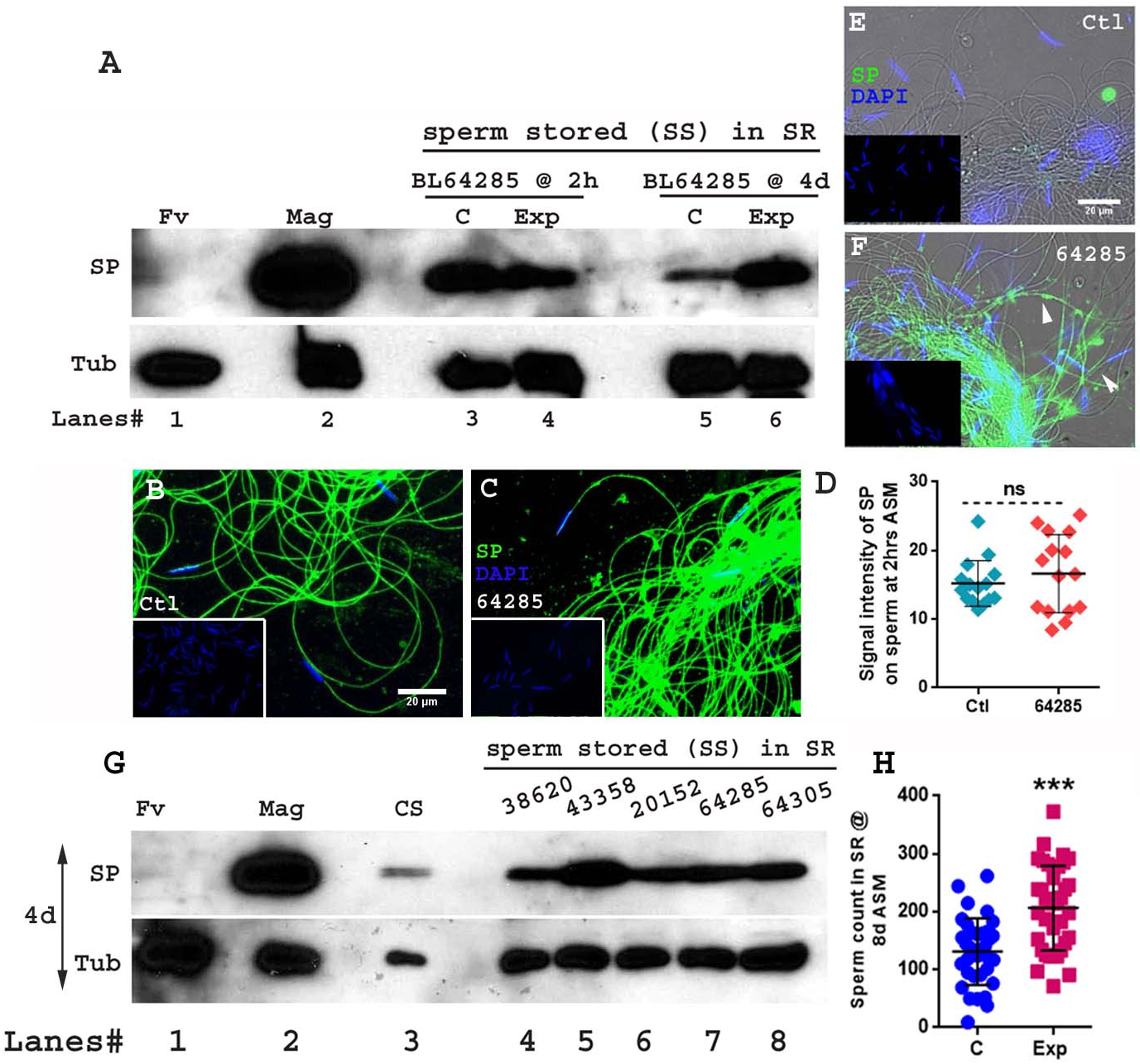
Loss of SSCs and/or parovaria in Hr39 mutant females does not inhibit the binding of SP to sperm but leads to retention of sperm and therefore elevated SP levels long term. **(A)** Western blot probed for SP. **Lanes# 1: Fv**, reproductive tract (RT) of 4 virgin females (negative control), **2: Mag**, 1 pair of male accessory glands (positive control), **3: B**,**C**, sperm dissected from SR of 30 genetically matched control (BL64285) females mated to wild type (CS) males at 2 hrs ASM, **4: Exp**, sperm dissected from SR of 30 mutant (BL64285) females mated to wild type (CS) males at 2 hrs ASM, **5: C**, sperm dissected from SR of 30 genetically matched control (BL64285) females mated to wild type (CS) males at 4 days ASM **6: Exp**, sperm dissected from SR of 30 mutant (BL64285) females mated to wild type (CS) males at 4 days ASM. Tubulin (Tub) served as loading control. Sperm samples isolated from the seminal receptacle of **(B)** matched-control females, **(C)** BL64285, Hr39 mutant females. The females were mated with CS males and frozen at 2 hrs ASM. **(D)** Relative signal intensity of SP bound to stored sperm in matched control (ctl) and BL64285 females at 2hrs ASM performed through immunofluorescence. Error bars show Mean±SE AU. Sperm samples isolated from the seminal receptacle of matched-control females **(E)**, BL64285, Hr39 mutant females (**F**). The females were mated with CS males and frozen at 4 days ASM. In all the immunofluorescence panels, sperm heads were stained with DAPI (blue) and anti-SP staining was visualized with Alexa fluor 488, staining the sperm tail (green) and sperm head (cyan; overlapping blue/green). The insets show the respective negative controls each panel. The bigger panels (E and F) in 4 days samples have transmitted light filter added to show the outline of sperm tail in the regions where SP was undetected (e.g, panel E). The white arrows indicate SP (green) on sperm tail. n=10; Bar = 20µm. **(G)** Western blot probed for SP at 4 days ASM. **Lanes# 1: Fv**, reproductive tract (RT) of 4 virgin females (negative control), **2: Mag**, 1 pair of male accessory glands (positive control), **3: CS**, sperm dissected from SR of 30 control (CS) females mated to wild type (CS) males, **4: 38620**, sperm dissected from SR of 30 Hr39 mutant (BL38620) females mated to wild type (CS) males, **5: 43358**, sperm dissected from SR of 30 Hr39 mutant (BL43358) females mated to wild type (CS) males, **6: 20152**, sperm dissected from SR of 30 Hr39 mutant (BL20152) females mated to wild type (CS) males, **7: 64285**, sperm dissected from SR of 30 Hr39 mutant (BL64285) females mated to wild type (CS) males, **8: 64305**, sperm dissected from SR of 30 Hr39 mutant (BL64305) females mated to wild type (CS) males. Tubulin (Tub) served as the loading control. **(H)** Sperm counts in SRs of genetically matched control (C; blue bar) and Hr39 mutant (Exp; pink bar) females from BL64285 stocks mated to control ProtB-eGFP males (with eGFP tagged sperm; p***=<0.001; n=15-20) and frozen at 8 days ASM. Error bars show Mean±SE AU (AU stands for arbitrary units).

Five day old Send1>CyO (control) and Send1>Rh1^G69D^ (experimental) females were mated to 3-day-old control (CS) males and the mated females were frozen at 2 hrs ASM. Sperm, dissected from seminal receptacles of the mated females, were assessed for the presence of SP by western blotting. We did not observe any striking difference in the levels of sperm-associated SP in experimental females relative to control females at 2hrs ASM (Fig. 4E lanes 3, 4). Similarly, there was no difference in amounts of LTR-SFPs CG1656, CG1652, Antr, or CG9997 associated with sperm isolated from experimental vs. control females at 2hr ASM (Fig 4E. lanes 3, 4). We also performed immunofluorescence on sperm dissected from the seminal receptacles of Send1>CyO (control) and Send1>Rh1^G69D^ (experimental) females to probe for the presence of SP. Consistent with the results of our western blots, we did not observe any striking difference in the intensity of anti-SP staining on sperm stored in experimental females (Fig. 4S B-B’ and E; Mean±SE=6.326±0.48 AU; p=0.8027) relative to those stored in control females (Fig. 4S A-A’ and E; 6.175±0.35 AU) at 2hrs ASM.

Likewise, five day old Hr39 mutant and control females were mated to 3 day old control (CS) males, and mated females were frozen at 2 hrs ASM. Genetically-matched controls were available for only one of the five available Hr39 mutant lines. We dissected sperm from seminal receptacles of genetically matched control (C; BL64285/CyO) and mutant (Exp) females from this line (BL64285 {Hr39[C105]}) [50] at 2hrs ASM and probed the samples for SP by western blotting and immunofluorescence. As we saw with Send1>Rh1^G69D^ vs. control females, we did not observe any striking difference in the levels of SP associated with sperm probed through western blotting (Fig. 5A lanes 3, 4; Fig. 5S3 A) or the signal intensity of anti-SP staining along the entire sperm performed through immunofluorescence in matched-control females (Fig. 5B and D; Mean±SE=15.21±0.8613 AU) when compared to mutant females at 2hrs ASM (Fig. 5C and D; 16.65±1.47 AU; p=0.4051). The other four available Hr39 mutant lines (BL38620 {Hr39[MI06174]} [51], BL43358 {Hr39[C277]} [52] and BL20152 {Hr39[EY04579]} [53]) did not have genetically-matched controls, or produced so few controls (BL64305/CyO {Hr39[c739]} [54]) that we could not perform experiments using them, so we used CS females as their controls. As we saw with BL64285 females and their controls, we detected similar levels of SP bound to sperm dissected from CS and these Hr39 mutant females (Fig. 5S2 A, lane 3, lanes 4-8; Fig. 5S2 B) at 2hrs ASM. We also detected signals for the LTR-SFPs CG1656, CG1652, Antr, and CG9997 in Hr39 mutant females at levels similar to those in CS females, at 2hr ASM (Fig. 5S3 C, lane 3 and lanes 4-8). The level of anti-SP staining visualized along the entire sperm through immunofluorescence also did not show any relative difference between CS females (control) (Fig. 5S2 C & H) and mutant females from the other four Hr39 lines (Fig. 5S2 D-H), consistent with what we observed in our western blots. Thus, loss of SSCs (and parovaria, in the case of Hr39 mutants) did not have an evident effect on the initial binding of SP or LTR-SFPs to sperm.

### 5. Loss of SSCs and parovaria in females affects the release of SP from stored sperm

Our Western blots, however consistently showed higher levels of sperm-associated SP at 4days ASM in Hr39 mutant females (BL 64285; Fig. 5A lane 6) relative to the levels in genetically matched-control females (Fig. 5A lane 5); this was particularly clear when SP levels were normalized with tubulin levels on the same blot (Fig. 5S3 B). We obtained analogous results showing higher levels of sperm-associated SP at 4 days ASM in the other four Hr39 mutant females (for which genetically matched control females were unavailable), relative to sperm-bound SP levels in CS females (Fig. 5G lanes, 3-8), following normalization of SP levels with tubulin (Fig. 5S4A).

Binding of SP to sperm or SP’s gradual cleavage from sperm are both essential for the efficient release of sperm from storage within the mated female [55]. To test whether the elevated levels of sperm-bound SP in Hr39 mutant females at 4d ASM was associated with increased retention of sperm in these females, we performed sperm counts. Control and Hr39 mutant females from the BL64285 stock were mated to ProtB-eGFP [56] males, and sperm stored in their seminal receptacle were counted at 8 days ASM. Mutant females (Fig. 5H, Exp) exhibited significantly higher sperm numbers relative to their genetically-matched controls (Fig. 5H, C; p**=<0.01). Consistently, mated Hr39 mutant females from the other four lines also showed significantly higher sperm counts, indicating poor release of stored sperm when compared to CS females (Fig. 5S3 D; p**=<0.01).

To distinguish whether the higher amounts of sperm-associated SP measured on Western blots was due to this higher retention of sperm or to impaired release of SP from sperm, we used immunofluorescence to examine levels of SP bound to sperm. We observed higher levels of sperm-bound SP in BL64285 Hr39 mutant females (Fig. 5F) than in matched control females (Fig. 5E); the latter’s SP levels at 4d ASM were below our detection limits. Similarly, anti-SP immunofluorescence was higher in females for the four other Hr39 mutants (Fig. 5S4 C-F) relative to levels in CS controls (Fig. 5S4 B); the latter again had SP levels at 4d ASM below our detection limits. We used Send1>Rh1^G69D^ flies, in an attempt to determine whether SSCs alone were responsible for effects on SP release. We saw no difference in SP levels bound to sperm dissected from Send1>CyO (control) and Send1>Rh1^G69D^ (experimental; SSCs ablated) females mated to CS (control) males (Fig. 4F, lane 3, 4 for 4 days ASM and 5, 6 for 8 days ASM), although signals were low in these experiments. Our results suggest that loss of some or all SSCs may not be sufficient to impair the release of SP from sperm

Our results indicate that the absence of SSCs and parovaria impairs the release of SP from sperm. Since the release of sperm from storage requires release of SP’s active region, by cleavage, from sperm, the lack of SP release in the absence of SSCs and parovaria result in more sperm being retained in females (and each of these sperm contains more bound SP than would normally be seen at those times). These data suggest that the protease responsible for cleaving SP’s active region from sperm [16] is provided by the parovaria and/or SSCs.

## Discussion

In addition to their crucial role in fertilization, sperm can have functions that modulate other aspects of reproduction. For example, Drosophila sperm can bind SP (and several other SFPs), causing SP’s retention in the female and allowing it to induce physiological and behavioral changes long-term [3,4,14,16]. Given that sperm thus can potentiate the effect of this male-derived protein, it is of interest to know whether the male, female, or both facilitate this binding. Previous studies showed that several SFPs are needed to bind SP to sperm, indicating male contributions to this phenomenon. Though several female proteins, to name a few, Fra mauro, Hadley, Esp, and Sex Peptide Receptor (SPR) have been identified that are needed for SP activity [24,28,29,31], none of these affected SP binding to sperm. Here, we show that female-derived factors are also necessary for SFPs, including SP, to bind to sperm. We report that levels of sperm-bound SFPs and SP are weak to undetectable in the male’s ejaculate, but increase once the ejaculate is within the FRT. Incubation experiments show that this increase is not simply due to time, but requires that the ejaculate be within the female, pointing to the need for female contributions to the SFP-sperm binding. We show that the FRT does not undergo drastic changes in pH following mating, and that secretions of SSCs and parovaria are also unlikely to be the critical female agents mediating initial SFP-sperm binding; we do however find a role for the latter in releasing SP from sperm. Our results suggest a molecular cooperation between male and female to bind SFPs to sperm.

### Levels of sperm-bound SP and LTR-SFPs increase within the mated female, though not all exhibit the same pattern (or kinetics)

SPs association with sperm is tightly regulated by a cascade of “LTR-SFPs” that prime sperm to bind SP [21,55]. Two LTR-SFPs, the CRISP CG17575, and the protease Seminase, do not themselves bind to sperm; instead, they facilitate the localization of other LTR-SFPs, and SP, to sperm and sperm storage organs [21,22]. Four other LTR-SFPs, the lectins CG1652, CG1656, the proteases CG9997, and the CRISP Antares (Antr) bind sperm transiently [14]. Like SP, CG9997 and Antr bind to both the head and tail of sperm, whereas CG1652 and CG1656 are detected only on the sperm tail [4,21].

Interestingly, we observed differences in the timing with which SP and individual LTR-SFPs associated with sperm. SP and two LTR-SFPs (CG1656 and Antr) were detected at low levels on sperm in ejaculate; the other LTR-SFPs were not detectable on sperm in ejaculate. Once ejaculate entered the female’s bursa, levels of SP, Antr, and CG1656 increased on sperm, but the other LTR-SFPs remained undetectable on sperm. Anti-SP staining on sperm in the bursa was spotty, rather than at higher and uniform staining seen in the seminal receptacle, indicating that its binding increased further once within the SR. In contrast, staining for Antr and CG1656 was already at maximal levels in the bursa and showed no further increase in the SR. Of the four remaining LTR-SFPs, sperm-binding by CG1652 and CG9997 was first detected in the SR (and seminase and CG17575 were undetectable on sperm even in the SR, consistent with previous reports [21,22]).

Several not-mutually-exclusive mechanisms could explain why different SFPs showed different kinetics of associating with sperm. First, it could be that some LTR-SFPs catalyze each other’s binding, and thus that some need to bind earlier than others. The earliest-binding LTR-SFPs may facilitate some SP binding, but full binding requires the full complement of LTR-SFPs on sperm. At least one LTR-SFP (CG9997) is post-translationally modified (cleaved) within the female. The role of this cleavage is unknown, but one could imagine that modifications of this sort could also affect a protein’s binding to sperm, or its ability to catalyze another protein’s sperm-binding.

Second, recently Wainwright et al., [57] showed that SP (at least) is transferred to females on large, neutral lipid-containing “microcarriers” that dissemble after entering the female reproductive tract, releasing their contents. It may be that the slow appearance of SP on sperm reflects its release by disassembly of these microcarriers; the timing that we observe is consistent with the timing of microcarrier dissociation reported by Wainright et al., [57]. A similar explanation could underlie the differences in sperm-binding kinetics of the LTR-SFPs, but it is not yet known whether they are transferred on microcarriers.

Third, recent results [32] show that as sperm transit through and remain in the FRT, female-derived proteins become associated with them. It is possible that some of these female proteins facilitate the binding of particular SFPs to sperm, and that the kinetics of each SFP’s association reflects the association of particular female proteins with sperm.

### The pH of the female reproductive tract is in the range of 6.0-7.4 and remains within this range in response to mating

The pH of the human FRT undergoes drastic change immediately after the deposition of semen. The low pH (4.3) of the vagina in humans is inhospitable to mammalian sperm. After mating, the vaginal pH rises to 7.2, creating an environment favorable for the viability and motility of sperm [43]. Given this finding, we tested whether similarly dramatic changes in FRT pH occurred in Drosophila post-mating, as this could potentially also be a factor promoting SFP-sperm binding. Using two sensors to examine the pH of the female reproductive tract before and after mating, we observed that the pH of the FRT was in the range of 6.0 to 7.4 and did not exhibit any drastic change outside of this range, post-mating. This is similar to reports in rodents (mice) which maintain their near-neutral vaginal pH after mating [38]. Because the available sensors (pHluorin R (ratiometric; [47]) and phluorin E (ecliptic; [44,45])) do not have the sensitivity to determine the pH more precisely than 6.0-7.4, we cannot determine whether mating shifts FRT pH within this range. Investigation of this will be an interesting area for future study, once new pH sensors with greater precision within this range become available.

### Partial or greater loss of spermathecal secretory cells or parovaria does not affect the initial association of SFPs with sperm, but affects the release of SP from sperm

When we disabled SSCs by driving expression of Rh1^G69D^ [48,49], or obtained full or partial loss of parovaria and SSCs with Hr39 mutants [41], we did not observe detectable differences in the initial amount (or distribution) of SFP-sperm association relative to controls, indicating that secretions from SSCs and/or parovaria do not play a major role in facilitating the initial binding of SFPs to sperm. However, Hr39 mutants differed from controls in the rate of release of SP from sperm; at 4d ASM, we observed higher retention of SP on sperm stored in Hr39 mutant females relative to levels seen in controls. The impaired release of SP from sperm that we observed in Hr39 females is expected to impair the rate by which these female release sperm from storage, as SP activity is needed for sperm release [55]. Consistent with this expectation, we observed that Hr39 mutant females from all five mutant lines showed significantly higher sperm counts in their SR at 8d ASM, verifying that they had poor release of stored sperm relative to control females. Our results may provide a mechanism for the observations in two previous studies [39,40] that secretions of the SSCs, or SSC and parovaria, are necessary for stored sperm to be efficiently used for fertilization.

The release of SP’s C-terminal active region from sperm occurs by proteolysis [16], but the source of the protease that accomplishes this has been a mystery. Our results suggest that this protease may be derived from the female, and specifically from her reproductive glands (or that its expression is regulated by the Hr39 transcription factor). That the female would provide the protease to release SP to sperm makes sense physiologically, in that SFPs (other than SP) do not persist in the female for more than one day post-mating, making it likely that a male-derived protease that could cleave SP would not remain in the female long enough to regulate SP cleavage (unless the protease is a sperm-protein). It also raises interesting evolutionary implications that the female would provide the activity that permits the active portion of SP to be released and function.

## Conclusion

Our findings, thus highlight that molecular contributions from both males and females are needed to facilitate association and/or dissociation of SFPs/SP to sperm and encourage future studies to identify the female candidates that mediate these molecular interactions between sexes.

## Methods

### Fly strains and crossing scheme

Flies used for ejaculate collections were derived from a cross between UAS-dTrpAI [58] and UAS-mCD8-Gfp; fru-GAL4(B)/MKRS [59]. The flies were a generous gift from the Baker lab (Janelia). Fru-GAL4>UAS-dTrpAI expel ejaculate after exposure to heat (29°) due to activation of the temperature-sensitive cation channel dTrpAI. Canton S (CS) females mated to CS males were used to collect sperm from bursa (35 min after the start of mating, ASM) and seminal receptacle (2hrs ASM). Ecliptic pHluorinE (UAS-MApHS; [45]) flies were a generous gift from the Han lab (Cornell University). Ratiometric pHluorinR, w[1118]; P{w[+mC]=UAS-pHMA}1.4, P{w[+mC]=UAS-pHMA}1.5A/CyO; TM2/TM6B, Tb[1] (BL44593; [47]) were obtained from the Bloomington Drosophila Stock Center (BDSC). All stocks not otherwise indicated were obtained from the BDSC. To disrupt/abolish the secretory units of the female reproductive tract, UAS-Rh1^G69D^ (flies were a generous gift from Dr. H.D. Ryoo; [48]) were crossed to Send1-GAL4; Gla/CyO (specific to spermathecae; kind gift of Dr.M. Siegal) and Oamb-GAL4 (specific to oviduct; kind gift of Dr. K. Han) flies to induce tissue specific generation of ER stress, and the ablation of secretory units (paraovaria and SSCs lining the spermathecal cap). We also used five-publicly available Hr39 mutant lines, y[1] w[*]; Mi{y[+mDint2]=MIC}Hr39[MI06174] (BL38620; [51]), w[*]; P{w[+mGS]=GSV1}Hr39[C277] (BL 43358; [52]), y[1] w[67c23]; P{y[+mDint2] w[+mC]=EPgy2}Hr39[EY04579] (BL20152; [53]), y[1] w[67c23]; Hr39[C105] (BL64285; [50]) and y[1] w[67c23]; P{w[+mW.hs]=GawB}Hr39[c739] P{w[+mC]=UAS-mCD8::GFP.L}LL5 (BL64305; [54]). Because genetically matched controls were either unavailable or sub-viable for 4/5 of the Hr39 mutant lines, we used CS females as relative controls. ProtB-eGFP males with Protamine B-eGFP tagged sperm heads were kindly gifted by the Pitnik lab [56]. All flies were reared under a 12:12h light-dark cycle at 22±1°C on standard yeast-glucose medium. Mating experiments were carried out by single-pair mating 3-5 day old unmated control males to 3-5 day old virgin females of the genotypes indicated in the text.

### Immunofluorescence

Immunostaining was performed to detect SP and LTR-SFPs binding to sperm as in [14,15]. Sperm isolated before (in male ejaculate) or after mating (in female bursa and seminal receptacle) were attached to poly-L-Lysine (Sigma) coated slides. Sperm isolated from male ejaculate were processed either immediately after exudation, or were incubated in 1X PBS for 2hrs after exudation from males. All the samples were processed according to the protocol of Ravi Ram and Wolfner [4] with minor modifications. Samples were blocked with 5% bovine serum albumin, BSA in 1X PBS for 30min. Subsequently, samples were incubated overnight in rabbit anti-SP(1:200), CG1656(1:100), CG1652(1:50), CG9997(1:50) [4,22,60] in 0.1% BSA at 4°C overnight. Samples were then washed in 1X PBS and incubated at room temperature for 2h in mouse anti-rabbit IgG coupled to Alexa fluor 488 (green; Invitrogen) at a concentration of 1:300 in 1x PBS at room temperature in the dark. Samples were then washed in PBS, incubated in 0.01% DAPI for 3 min at room temperature in the dark, rewashed and mounted using antifade (CitiFluor mountant solution; EMS). The fluorescence was visualized under an Echo-Revolve fluorescence microscope at a magnification of 20X. The intensity of anti-SFP immunofluorescence on sperm tails was quantified using Image J software (National Institute of Health, Bethesda, USA). The difference in the fluorescence intensity of anti-SFP on sperm tails between ejaculate, bursa and SR samples were statistically analyzed using one-way analysis of variance (ANOVA), followed by Dunnet’s multiple comparison tests. A minimum of three independent immunostaining batches, each with a minimum sample size of 10, were analyzed for each group.

### Visualizing the pH in the female reproductive tract

Ten virgin females per batch expressing ecliptic pHluorin-Oamb>MApHS, Send1>MApHS and ratiometric pHluorin-Oamb>pHMA, Send1>pHMA were dissected to detect the presence of pHluorin signals. Another ten of these females were mated to CS males and dissected at 35 minutes ASM to detect and analyze the pHluorin signals (relative to virgin females). Whole female reproductive tracts were dissected in 1X PBS on a slide. The tissues were fixed with 4% paraformaldehyde (PFA) for 15 min. Samples were then washed in 1X PBS, incubated in 0.01% DAPI for 3 min at room temperature in the dark, rewashed and mounted using antifade (CitiFluor mountant solution; EMS). Images were captured through an Echo-Revolve fluorescence microscope at a magnification of 20X. A minimum of three independent batches, with a minimum sample size of 10 per batch, were analyzed for each group.

### Efficacy of ablation of SSCs

The reproductive tracts from Send1>Rh1^G69D^ and Hr39 mutant females were dissected and analyzed to detect the presence of ablated SSCs, if any. Whole female reproductive tracts were dissected in 1X PBS on a slide. The tissues were fixed with 4% PFA, further processed, mounted and imaged the same way as described above for pH change visualization in the female reproductive tracts. A minimum of two independent batches, with a minimum sample size of five per batch, were analyzed for each group.

### Sample preparation and Western blotting

To determine the binding of SP and LTR-SFPs to sperm and persistence of sperm bound SP long-term, sperm stored (SS) in the seminal receptacle of females (of the indicated genotype) mated to control males were dissected. The dissected tissues (SS, n=30) were suspended in 5µl of homogenization buffer (5% 1M Tris; pH 6.8, 2% 0.5M EDTA) and processed further according to the protocol of Ravi Ram and Wolfner [4]. Proteins from stored sperm were then resolved on 12% polyacrylamide SDS gel and processed further for western blotting. Affinity purified rabbit antibodies against SP(1:2000), CG1656(1:1000), CG1652(1:500), antares(1:500), CG9997(1:1000), CG17575(1:1000), seminase(1:1000) [4,14,22] and mouse antibody against tubulin (as a loading control; 1:3500) were used as primary antibodies. HRP conjugated secondary anti-rabbit and anti-mouse antibodies (Jackson Research) were used for detection of SFPs at a concentration of 1:2000. The levels of SP were normalized with tubulin of respective lanes using Quantity One software.

### Sperm release from sperm storage organs in females

To study the sperm utilization and release, Hr39 mutant females mated to ProtB-eGFP (control) males were frozen at 8d ASM for sperm counts. Subsequently, seminal receptacles of mated females were dissected and eGFP sperm were counted (at a total magnification of 20X, with FITC filter on an Echo-Revolve microscope). Mature sperm in the seminal receptacles of mated females were counted twice and groups were blinded to ensure reproducibility and avoid bias [61]. The percent repeatability was 88-92%. Assays were repeated twice, with two technical replicates. Differences in the sperm counts between groups were analyzed statistically through one-way ANOVA followed by Tukey’s multiple comparison tests. Each group contained a minimum sample size of 15-25.

## List of abbreviations

SFP: seminal fluid protein
SP: Sex Peptide
LTR-SFPs: long-term response SFPs
FRT: female reproductive tract
SSC: spermathecal secretory cell
ASM: after the start of mating
MApHS: membrane-associated pH sensor
AU: arbitrary unit

## Declarations

### Ethics approval and consent

Not applicable: there were no human subjects or vertebrate animals used in this research.

### Data availability

All data generated or analysed during this study are included in this manuscript and its supplementary information files.

### Consent for publication

Not applicable.

### Author contributions

S.M, A.S and M.F.W designed the experiments; S.M and A.S carried out the experiments; S.M and M.F.W analyzed the results. S.M and M.F.W wrote and revised the manuscript.

## Acknowledgements

We are grateful to the NIH for grant R37-HD038921 (to MFW) for support. We thank Dr. Chun Han for the pH-sensor stocks and suggestions, the Baker lab (Janelia) for UAS-dTrpAI and UAS-mCD8-Gfp; fru-GAL4(B)/MKRS stocks, Dr. H.D Ryoo for the UAS Rh1^G69D^ stocks and Bloomington Drosophila Stock Center for Hr39 mutant stocks. We are grateful to I.A. Amaro, D.S. Chen, Y. Hafezi, J. Thomalla, S. Allen and M. Yang for helpful suggestions or for comments on the manuscript.

