## Supplemental Material for "Female factors are important for the seminal Sex Peptide’s association with sperm in mated D. melanogaster"

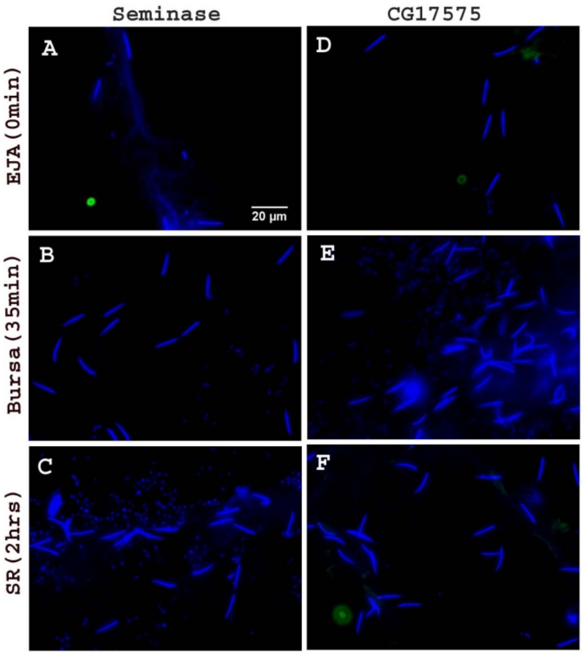


**Figure 2S**. **CG17575 and Seminase do not associate with sperm at any stage from male ejaculate to female storage.** Pre-mating ejaculate samples were collected from Fru>dTRPA1 males exposed to high temperature. Post mating sperm samples were isolated from mated females. Wild type (CS) females were mated to a wildtype (CS) male and frozen at 35 min (sperm in bursa) and 2 hrs (sperm stored in seminal receptacle) ASM. Sperm heads were stained with DAPI (blue) and seminase and CG17575 were visualized with Alexa fluor 488 (green). Sperm isolated from male ejaculate **(A and D)** probed for seminase and CG17575, respectively. Sperm isolated from female bursa at 35 min ASM **(B and E)** probed for seminase and CG17575, respectively. Sperm isolated from female seminal receptacle at 2hrs ASM **(C and F)** probed for seminase and CG17575, respectively. (n=10; Bar = 20μm)

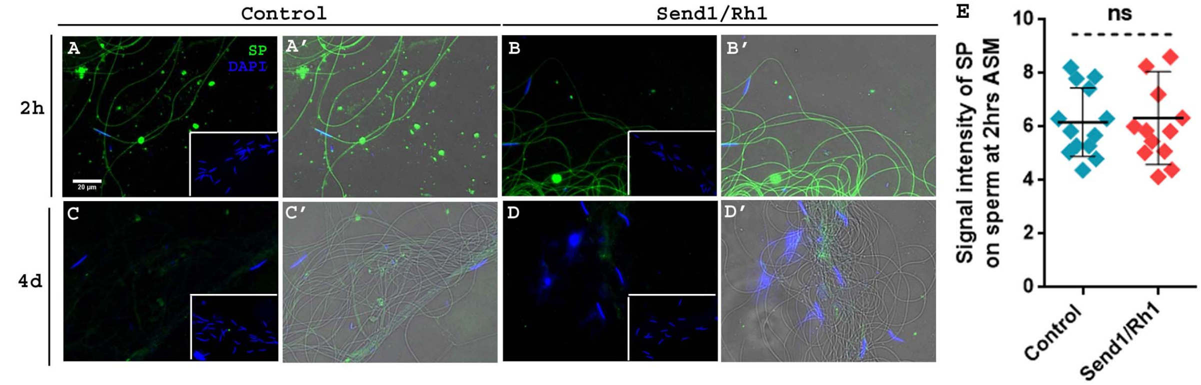


**Figure 4S1. Anti-SP staining on sperm dissected from Send1> Rh1 females (with ablated SSCs) do not show any difference in SP levels when compared to their levels in matched-control females at 2hrs or 4 days ASM.** Sperm samples isolated from the seminal receptacle of *Send1>CyO* (Control) females **(A-A')** and *Send1>Rh1* (experimental) females **(B-B')** mated with CS males and frozen at 2 hrs ASM. Sperm samples isolated from the seminal receptacle of *Send1>CyO* (Control) females **(C-C')** and *Send1>Rh1* (experimental) females **(D-D')** mated with CS males and frozen at 4days ASM. Sperm heads were stained with DAPI (blue) and anti-SP staining was visualized with Alexa fluor 488, staining the sperm tail (green) and sperm head (cyan; overlapping blue/green). Insets show the respective negative controls for each panel, with only secondary antibody (anti-rabbit, Alexa fluor 488) and no primary antibody (anti-SP) incubation. Panels **A', B', C', D'** have an added transmitted light filter to show the outlines of sperm tails in the regions where SP was undetected (e.g, panel C and D), n=10; Bar = 20µm **(E)** Relative signal intensity of SP bound to stored sperm in *Send1>CyO* (control) and *Send1>Rh1* females at 2hrs ASM; ns= non significant; error bars show Mean±SE AU (AU stands for arbitrary units).

1. ***Ablation of spermathecal secretory cells (SSCs) in the female reproductive tract varied in degree and penetrance of the phenotype across the lines tested***

We ablated the SSCs that line the spermathecal cap by driving the expression of misfolded protein $Rh^{1G69D}$ in these cells [1, 2]. Expression of $Rh^{1G69D}$ induces excessive ER stress in the targeted tissues, disrupting protein synthesis/secretion by the cells. We used *Send1-Gal4* to drive the expression of $Rh^{1G69D}$ in the female reproductive tract. *Send1>Rh*$^{1G69D}$ females had ablated SSCs, but the penetrance of the phenotype was not complete: some of the mosaic females (3/5) had only one of the two spermathecae lacking SSCs and the other spermatheca still had numerous SSCs (Fig. 4 C and D). Control *Send1>CyO* females had normal numbers of SSCs around their spermathecal caps (Fig. 4 A and B; DAPI stained).

As an alternative way to ablate SSCs, we used mutants of the nuclear hormone receptor Hr39 [3, 4], which is needed for the formation of secretory units in the female reproductive tract. Loss of expression of Hr39 affects the development of SSCs and parovaria. We focused on *Hr39* mutants that exhibit more stringent phenotypes for SSC ablation. We tested five different *Hr39* mutants- BL38620 (*Hr39[MI06174]*), BL43358 (*Hr39[C277]*), BL20152 (*Hr39[EY04579]*), BL64285 (*Hr39[C105]*) and BL64305 (*Hr39[c739]*) to determine the extent of loss of SSCs in mutant females from each stock relative to control females. Ablated SSCs were observed around the spermathecal caps in the reproductive tracts of all the five Hr39 mutant lines, but again, the penetrance of phenotype varied in SSC numbers (or size). As expected, CS females had large, regular SSCs (approx 60-80 cells) lining both spermathecal caps (Fig. 5S1 A and B; DAPI stained). The mutant BL 38620 females gave us the same pattern of SSC ablation as was observed in *Send1>Rh1*$^{G69D}$ females where some females lacked SSCs in both spermathecae (3/5), while other females (2/5) were mosaic, with only one of the two spermathecae lacking SSCs (Fig. 5S1 C and D; DAPI stained). Mutant females in stocks BL43358 (Fig. 5S1 E and F;

DAPI stained) and BL 64285 (Fig. 5S1 I and J; DAPI stained) had completely ablated SSCs in some females (4/5) and extremely reduced numbers and sizes of the SSCs lining the spermathecal cap, in others (1/5). The other two stocks, BL 20152 (Fig. 5S1 G and H; DAPI stained) and BL64305 (Fig. 5S1 K and L; DAPI stained) had females with almost complete ablation of SSCs from both the spermathecae, except that 3-4 SSCs could still be seen in some (1/5) of the females.

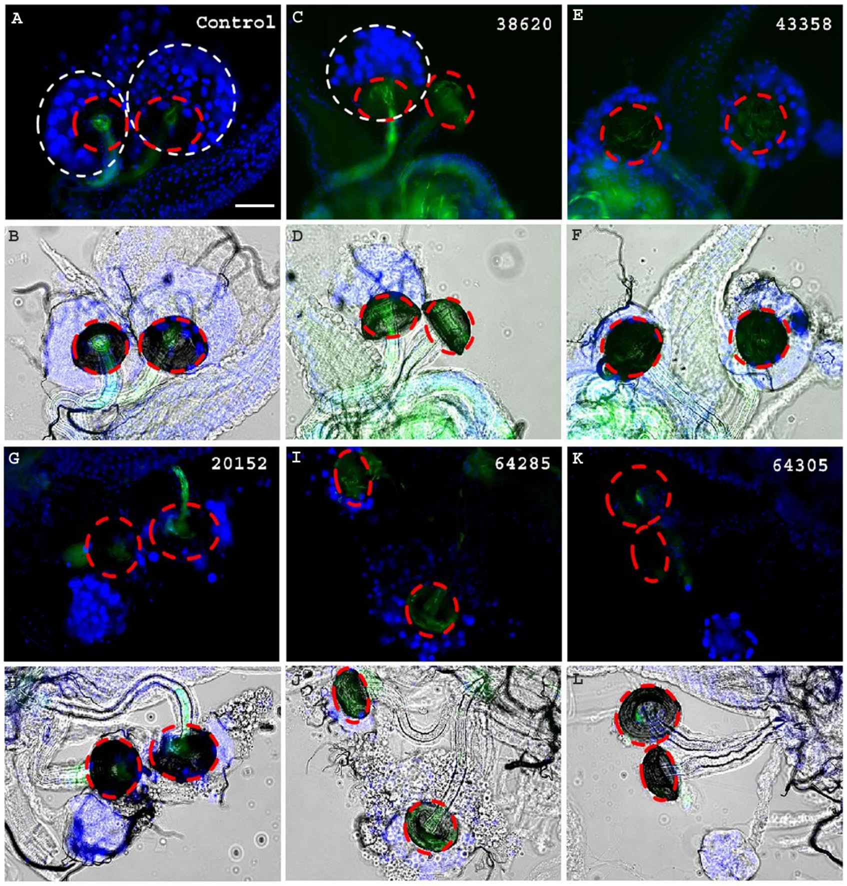


**Figure 5S1.** ***Hr39*** **mutant females show either completely ablated or extremely reduced SSC numbers.** Control or mutant females were mated with *ProtB-eGFP* males (eGFP-tagged sperm; green). SSCs, marked with DAPI stained nuclei (cells enclosed in white dotted circle) lining the spermathecal cap (red dotted circle). **(A-B)** Control (CS) females show normal bunch of SSCs around both the spermathecal caps. Completely ablated SSCs or SSCs reduced in cell size or number were observed in the *Hr39* mutant lines **(C-D)** BL38620, **(E-F)** BL 43358, **(G-H)** BL20152, **(I-J)** BL 64285, **(K-L)** BL 64305. n=5; Bar = 20μm.

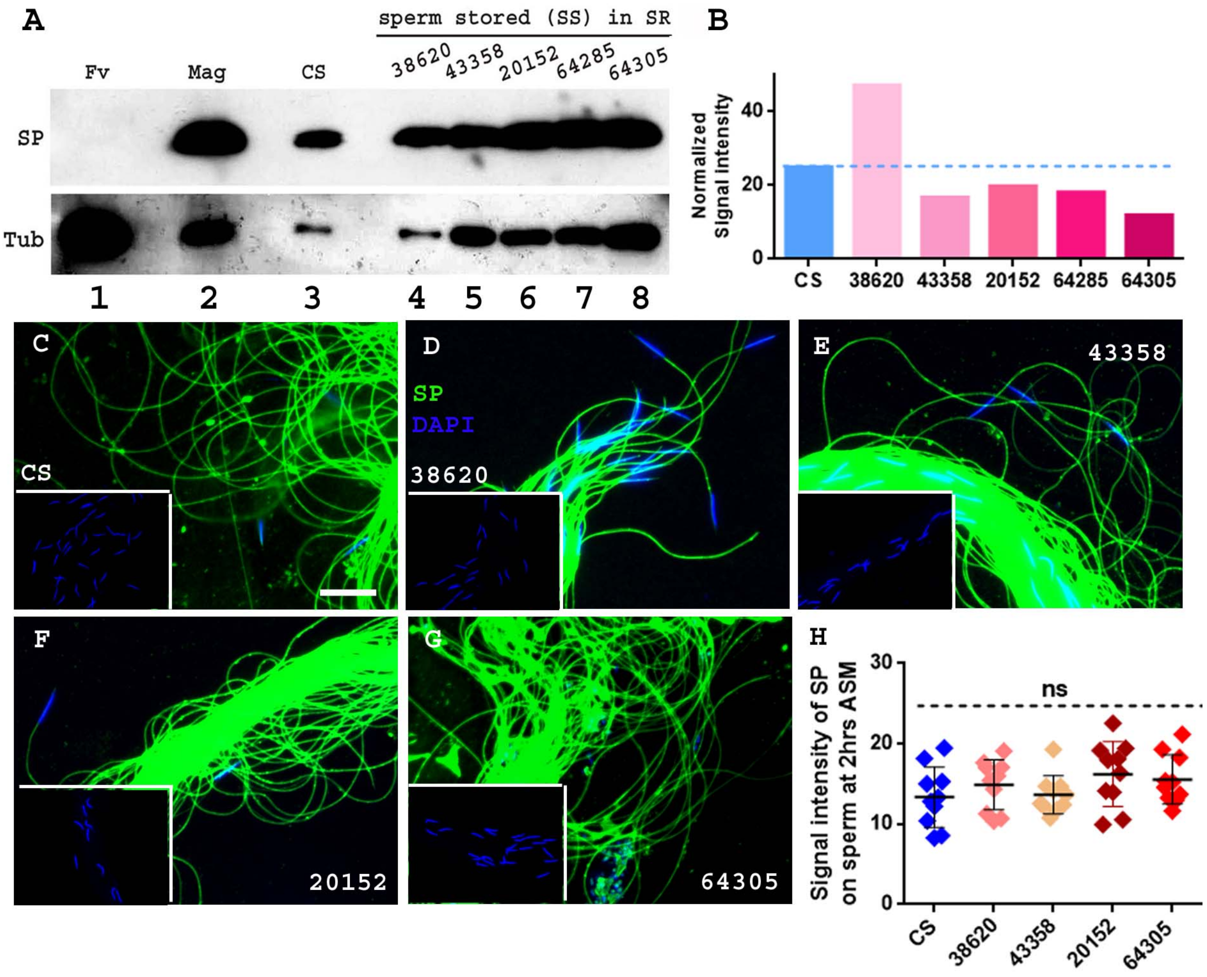


**Figure 5S2. Initial SP binding with sperm dissected from *Hr39* mutant females (with ablated SSCs) do not show any difference in SP levels when compared to their levels in CS females at 2hrs ASM. (A)** Western blot probed for SP at 2hrs ASM. **Lanes# 1: Fv,** reproductive tract (RT) of 4 virgin females (negative control), **2: Mag**, 1 pair of male accessory glands (positive control), **3: CS**, sperm dissected from SR of 30 control (CS) females mated to wild type (CS) males, **4: 38620,** sperm dissected from SR of 30 *Hr39* mutant (BL38620) females mated to wild type (CS) males, **5: 43358**, sperm dissected from SR of 30 *Hr39* mutant (BL43358) females mated to wild type (CS) males, **6: 20152**, sperm dissected from SR of 30 *Hr39* mutant (BL20152) females mated to wild type (CS) males, **7: 64285**, sperm dissected from SR of 30 *Hr39* mutant (BL64285) females mated to wild type (CS) males, **8: 64305,** sperm dissected from SR of 30 *Hr39* mutant (BL64305) females mated to wild type (CS) males. Tubulin (Tub) served as the loading control. **(B)** Graphical representation of the normalized levels of sperm bound SP in *Hr39* mutant (red bars) females from all the five stocks relative to CS females (blue bar & blue dotted line) at 2hrs ASM, as seen on one of three replicate Western blots; the other two blots showed similar results. Sperm samples isolated from the seminal receptacle of CS (control) females **(C)**, and other four *Hr39* mutant females, BL38620 **(D)**, BL43358 **(E),** BL20152 **(F)**, BL64305 **(G).** The females were mated with CS males and frozen at 2 hrs ASM. Sperm heads were stained with DAPI (blue) and anti-SP staining was visualized with

Alexa fluor 488, staining the sperm tail (green) and sperm head (cyan; overlapping blue/green). Insets show the respective negative controls for each panel, with only secondary antibody (anti-rabbit, Alexa fluor 488) and no primary antibody (anti-SP) incubation, n=10; Bar = 20µm **(H)** Graphical representation of relative signal intensity of SP bound to stored sperm in CS females (control) and other four Hr39 females, BL38620, BL43358, BL20152 and BL64305 at 2hrs ASM; ns= non-significant. Error bars show Mean±SE AU.

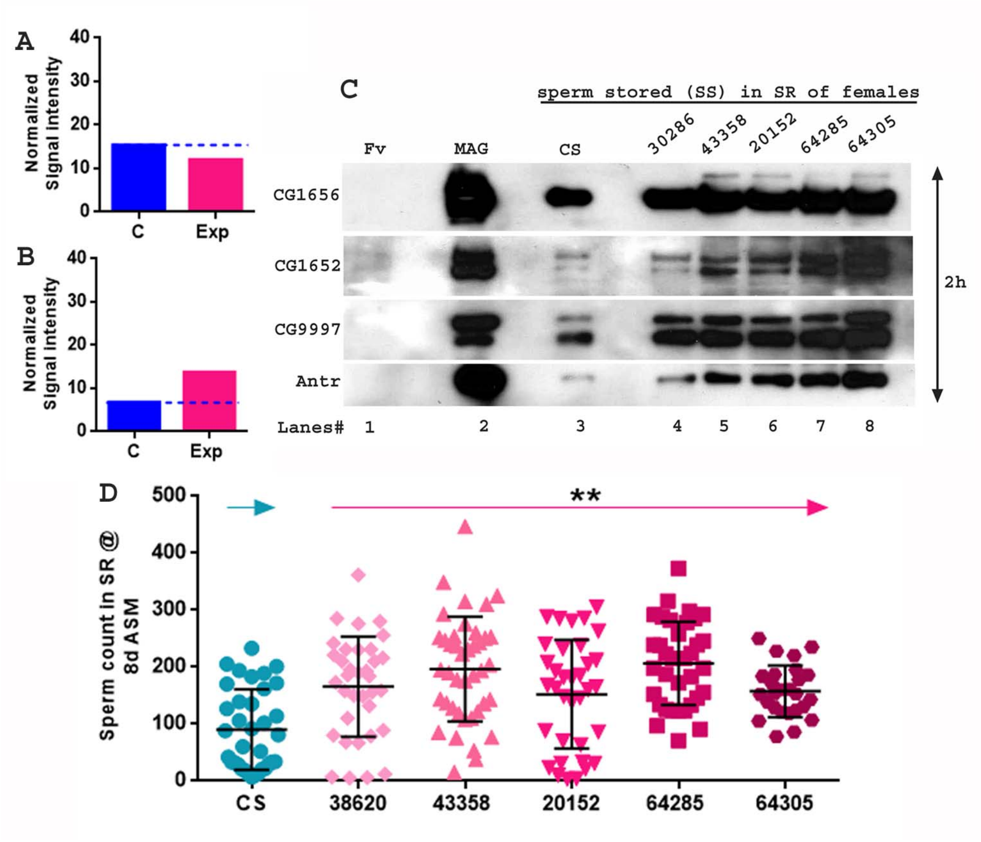


**Figure 5S3. *Hr39* mutant females have normal binding of LTR-SFPs to sperm but excessively retain sperm in storage. (A)** Graphical representation of the normalized levels of sperm bound SP in *Hr39* mutant (exp; pink bar) females relative to genetically matched control females (C; dark blue bar & dotted line) from the stock BL64285, at 2hrs ASM. **(B)** Graphical representation of the normalized levels of sperm bound SP in *Hr39* mutant (exp; pink bar) females relative to genetically matched control females (C; dark blue bar & dotted line) from the stock BL64285, at 4 days ASM **(C)** Western blot probed for LTR-SFPs, at 2hrs ASM. **Lanes# 1: Fv,** reproductive tract (RT) of 4 virgin females (negative control), **2: Mag**, 1 pair of male accessory glands (positive control), **3: CS**, sperm dissected from SR of 30 control (CS) females mated to wild type (CS) males, **4: 38620,** sperm dissected from SR of 30 *Hr39* mutant (BL38620) females mated to wild type (CS) males, **5: 43358**, sperm dissected from SR of 30 *Hr39* mutant (BL43358) females mated to wild type (CS) males, **6: 20152**, sperm dissected from SR of 30 *Hr39* mutant (BL20152) females mated to wild type (CS) males, **7: 64285**, sperm dissected from SR of 30 *Hr39* mutant (BL64285) females mated to wild type (CS) males, **8:**

**64305,** sperm dissected from SR of 30 *Hr39* mutant (BL64305) females mated to wild type (CS) males. Lanes were probed for LTR-SFPs , CG1656, CG1652, Antares and CG9997 as described in the text. **(D)** Graphical representation of sperm counts in SRs of CS and *Hr39* females from all the five mutant stocks, mated to control *ProtB-eGFP* males (with eGFP tagged sperm; green; error bars show mean±SE; p**=<0.01; n=15-20) and frozen at 8 days ASM.

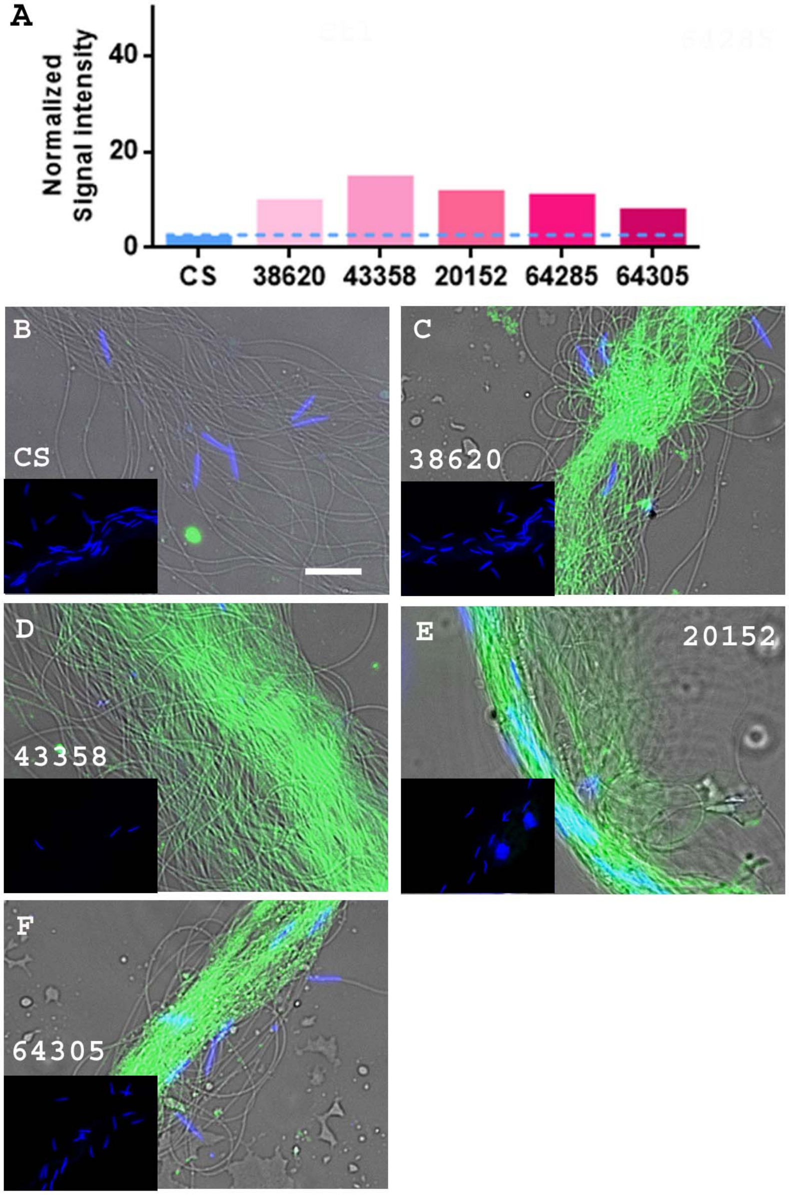


**Figure 5S4. SP levels on sperm after normalization on western blot and anti-SP staining on sperm dissected from *Hr39* mutant females (with ablated SSCs) show higher levels of SP levels when compared to their levels in matched-control or CS females at 4 days ASM. (A)** Graphical representation of the normalized levels of sperm bound SP in *Hr39* mutant (red bars)

females from all the five stocks relative to CS females (blue bar & dotted line) at 4 days ASM, as seen on one of three replicate Western blots; the other two blots showed similar results. Sperm samples isolated from the seminal receptacle of **(B)** CS (control) females and the other four Hr39 mutant females, **(C)** BL38620, **(D)** BL43358, **(E)** BL20152, **(F)** BL64305**.** The females were mated with CS males and frozen at 4 days ASM. Sperm heads were stained with DAPI (blue) and anti-SP staining was visualized with Alexa fluor 488, staining the sperm tail (green) and sperm head (cyan; overlapping blue/green). The insets show the respective negative controls for their panels. The larger panels have transmitted light filter added to show the outline of sperm tail in the regions where SP was undetected (e.g, panel B); n=10; Bar = 20μm.


1. Ryoo HD, Domingos PM, Kang MJ, Steller H. Unfolded protein response in a Drosophila model for retinal degeneration. EMBO J. 2007;26:242–52. doi:10.1038/SJ.EMBOJ.7601477.

2. Chow CY, Avila FW, Clark AG, Wolfner MF. Induction of excessive endoplasmic reticulum stress in the Drosophila male accessory gland results in infertility. PLoS One. 2015;10. doi:10.1371/JOURNAL.PONE.0119386.

3. Sun J, Spradling AC. NR5A nuclear receptor Hr39 controls three-cell secretory unit formation in Drosophila female reproductive glands. Curr Biol. 2012;22:862–71. doi:10.1016/J.CUB.2012.03.059.

4. Schnakenberg SL, Matias WR, Siegal ML. Sperm-storage defects and live birth in Drosophila females lacking spermathecal secretory cells. PLoS Biol. 2011;9. doi:10.1371/JOURNAL.PBIO.1001192.
